# Cannabis Expression Atlas: a comprehensive resource for integrative analysis of *Cannabis sativa* L. gene expression

**DOI:** 10.1101/2024.09.27.615413

**Authors:** Kevelin Barbosa-Xavier, Francisnei Pedrosa-Silva, Fabricio Almeida-Silva, Thiago M. Venancio

## Abstract

*Cannabis sativa* L., a plant originating from Central Asia, is a versatile crop with applications spanning textiles, construction, pharmaceuticals, and food products. This study aimed to compile and analyze publicly available Cannabis RNA-Seq data and develop an integrated database tool to help advance Cannabis research in various topics such as fiber production, cannabinoid biosynthesis, sex determination, and plant development. We identified 515 publicly available RNA-Seq samples that, after stringent quality control, resulted in a high-quality dataset of 394 samples. Utilizing the Jamaican Lion genome as reference, we constructed a comprehensive database and developed the Cannabis Expression Atlas (https://cannatlas.venanciogroup.uenf.br/), a web application for visualization of gene expression, annotation, and functional classification. Key findings include the quantification of 27,640 Cannabis genes and their classification into seven expression categories: not-expressed, low-expressed, housekeeping, tissue-specific, group-enriched, mixed, and expressed-in-all tissues. The study revealed substantial variability and coherence in gene expression across different tissues and chemotypes. We found 2,382 tissue-specific genes, including 177 transcription factors.. The Cannabis Expression Atlas constitutes a valuable tool for exploring gene expression patterns and offers insights into Cannabis biology, supporting research in plant breeding, genetic engineering, biochemistry, and functional genomics.

## 1 Introduction

*Cannabis sativa* L., commonly referred to as Cannabis, hemp, marijuana, or pot, originates from Central Asia and stands among the most versatile crops, serving as a fundamental resource for textiles, paper, construction materials, pharmaceuticals, and food products (Gao *et al*., 2020). With a cultivation history spanning over 10,000 years, Cannabis has been integral to textile production in China for over 6,000 years (Hussain *et al*., 2021). Evidence from Central Asia indicates that the utilization of drug-type genotypes, particularly for medicinal and ritualistic purposes, dates back to over 2,700 years (Ren *et al*., 2019).

Currently, the global market for hemp products generates over 100 million US dollars (US$) annually (United Nations, 2024). From 2019 to 2022, the global export of hemp seeds averaged 122 million US$, with Canada being the largest exporter, accounting for 46.54% in 2022. By 2022, 41 countries were exporting hemp seeds or related products (United Nations, 2024). Meanwhile, hemp fibers averaged 14.4 million US$ in trade between 2019 and 2022. During the same period, an average of 40 countries imported hemp fiber products, and about 25 countries exported them, with European countries dominating the market (United Nations, 2024).

Cannabis has a diploid genome (2n = 20) comprising nine pairs of autosomes and one pair of sex chromosomes, rendering it a dioecious species wherein male and female plants exhibit XY and XX sex chromosomes, respectively (Grassa *et al*., 2018). The Cannabis Y chromosome has an entire heterochromatic arm and a large pseudoautosomal region on the other arm indicating that some genes have homologs at the X chromosome and have autosomal behavior (Divashuk *et al*., 2014).

The primary feature that makes Cannabis widely utilized for medicinal, ceremonial, and adult purposes is its diverse cannabinoid profile (ElSohly and Slade, 2005). Several cannabinoid compounds have been identified, with Tetrahydrocannabinol (THC) and Cannabidiol (CBD) emerging as the most widely recognized. Based on their cannabinoid profiles, Cannabis plants can be categorized into different chemotypes (de Meijer *et al*., 2003). Type I, commonly known as marijuana, primarily contains tetrahydrocannabinolic acid (THCA) as the predominant compound. Type II plants exhibit similar levels of THCA and cannabidiolic acid (CBDA). Finally, type III plants, also known as hemp, feature CBDA as the primary compound. Cannabis plants synthesize THCA and CBDA, which are decarboxylated to THC and CBD, respectively. Legislation in various countries often relies on THC concentration to distinguish between marijuana and hemp. Typically, plants containing more than 0.3% THC are classified as marijuana, while those with less than 0.3% THC are considered hemp (Stout *et al*., 2012). Scientific progress in Cannabis research, including DNA and RNA sequencing, has greatly improved our comprehension of the molecular mechanisms underlying fiber production, cannabinoid biosynthesis, sex determination, and plant development (Guerriero *et al*., 2017; McKernan *et al*., 2020; Adal *et al*., 2021; Tang *et al*., 2023). Over the past 13 years, several research groups have generated a substantial volume of Cannabis RNA-Seq data (Bakel *et al*., 2011; Guerriero *et al*., 2017; Prentout *et al*., 2020). Nevertheless, there is a lack of systematic integrative analyses of these datasets, which often require specialized personnel and computational resources beyond the means of most research groups. Moreover, databases with visualization tools that integrate gene expression data from multiple chemotypes and tissues are crucial to accelerate research projects, and empower Cannabis researchers worldwide.

In this study, we performed an extensive analysis of all publicly available Cannabis RNA-Seq data, resulting in a curated dataset of 394 high-quality samples. We estimated transcriptional abundances of 27,640 genes, including 1,465 transcription factors (TFs). We further classified these genes into seven expression categories, including tissue-specific and housekeeping, providing valuable insights into Cannabis functional genomics. To enhance accessibility and usability, we developed a user-friendly web application, the Cannabis Expression Atlas (Cannatlas; https://cannatlas.venanciogroup.uenf.br/), that enables researchers to explore gene expression and molecular biology of Cannabis, supporting plant breeding, genetic engineering, and medicinal research.

## 2 Materials and Methods

### Data acquisition, quality check, and transcript abundance estimation

We identified the available Cannabis RNA-Seq data on the Sequence Read Archive (SRA) database using the search parameters “Cannabis sativa”[organism] AND “rna-seq”[strategy]. We used the R package ‘bears’ (Almeida-Silva, Pedrosa-Silva and Venancio, 2023) to obtain sample metadata with the create_sample_info() function. The FASTQ files were downloaded from the European Nucleotide Archive’s mirror of SRA using the download_from_ena() function, and file integrity was verified with the check_md5() function. Adapters and low quality bases were removed using FASTP (Chen *et al*., 2018). At this stage, we excluded samples with mean read length of less than 40 or a Q20 rate below 80% after filtering (Almeida-Silva, Pedrosa-Silva and Venancio, 2023).

The most commonly used Cannabis reference genomes are Finola, Purple Kush, and Cs10 (Grassa *et al*., 2018; Laverty *et al*., 2019), which belong to type I and III female plants. This has two implications: i. they do not have male-specific genome segments, and ii. type I plants do not have a functional CBDAS gene, while type III plants do not have a functional THCAS gene (McKernan *et al*., 2020). McKernan *et al*. (2020) reported the genome of a type II female and male plant, the Jamaican Lion Mother (JL mother) and the Jamaican Lion Father (JL father), and found male-specific contigs, making this the most complete Cannabis genome to date. The JL mother and JL father are available in GenBank under accession numbers GCA_012923435.1 and GCA_013030025.1, respectively. We used as reference the JL mother genome, complemented with the Y contigs from the JL father (McKernan *et al*., 2020) as a reference in our atlas. This genome was used to build a reference transcriptome with the R packages GenomicFeatures and Rsamtools (Lawrence *et al*., 2013; Martin Morgan *et al*., 2023).

We estimated gene-level transcript abundances with ‘salmon’ (Patro *et al*., 2017), which performs pseudo-mapping to the reference transcriptome followed by quantification. At this stage, we excluded samples with less than 50% mapped reads (Almeida-Silva, Pedrosa-Silva and Venancio, 2023). Additionally, genes with less than 1 transcript per million (TPM) in all samples were classified as not-expressed and not used in the downstream analysis.

### Functional gene annotation

Functional gene annotation was conducted using an integration of three database tools: InterProScan 5, UniProt - IDmapping, and KEGG - GhostKOALA (Jones *et al*., 2014; Kanehisa, Sato and Morishima, 2016; The UniProt Consortium, 2023). These tools enabled us to construct an annotation matrix with the following information: Gene ID, Protein ID, Entry name (UniProt), Protein name (UniProt), Gene Ontology (UniProt), Accession (InterPro), Description (InterPro), Gene Ontology (InterPro), Pathway annotations (InterPro), Entry (KEGG), Definition (KEGG), Pathway (KEGG), and Module (KEGG). TF prediction was conducted using PlantTFDB v5.0 (Tian *et al*., 2020).

### Dimensionality reduction

Dimensionality reduction methods were used to evaluate sample clustering patterns. The ‘scran’ package (Lun, McCarthy and Marioni, 2016) was used to model the mean-variance relationship in a matrix of log-transformed counts normalized by library size, followed by the identification of the top 5000 genes with the highest expression variability. Next, we performed a principal component analysis (PCA) and used the elbow point statistic to extract the top principal components. We employed the t-stochastic neighbor embedding (t-SNE) (Van Der Maaten and Hinton, 2008) dimensionality reduction method to compare the data distribution in a low-dimensional space. We tested 6 different perplexity values (10, 20, 30, 40, 50, and 60) and selected 30 as the optimal value based on visual inspection.

### Gene classification by expression category

We identified tissue-specific genes for tissues with at least 10 samples (leaf, trichome, female flowers, roots, hypocotyls, stem, male flowers, seeds, and bast fibres). Although induced male flowers had only 6 samples, we kept them to identify tissue-specific genes. Samples from mixed tissues were excluded. We obtained the median TPM expression of each gene in each tissue, as previously described by (Machado *et al*., 2020; Almeida-Silva, Pedrosa-Silva and Venancio, 2023). Genes with median TPM of at least 5 in a tissue were classified using the ‘TissueEnrich’ R package (Jain and Tuteja, 2019) and had their tissue specificity index TAU calculated using the following formula:

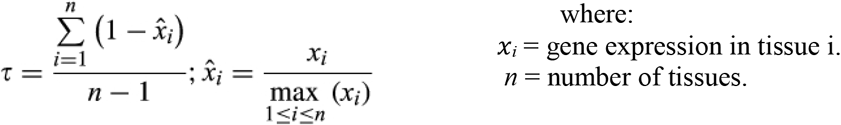

Genes identified as “Tissue-Enriched” or “Tissue-Enhanced” by the teGeneRetrival() function of the ‘TissueEnrich’ package and with TAU ≥ 0.8 were classified as “tissue-specific” genes. Furthermore, ‘TissueEnrich’ also returns other three gene expression categories, the “Group-Enriched”, “Mixed”, and “Expressed-in-all”.

### Identification of housekeeping genes

To consistently identify housekeeping genes, we utilized the methods previously employed in the Soybean Expression Atlas developed by our group (Machado *et al*., 2020; Almeida-Silva, Pedrosa-Silva and Venancio, 2023):

- Selected the genes with TPM ≥ 5 in at least one sample;
- Selected genes that are expressed in all samples;
- Obtained the median TPM of each gene across all samples;
- Computed the standard deviation (sd) and the coefficient of variation (CoV) of gene expression;
- Computed the maximum fold change (MFC) by determining the largest and smallest TPM value;
- The MFC-CoV score was calculated by multiplying the MFC with the CoV;
- Identified the first quartile;
- Identified the Tau index for the housekeeping putative genes.

Genes with MFC-CoV score within the first quartile and with a Tau index ≤ 0.4 were classified as housekeeping genes.

### Data visualization

Data visualization was mainly performed using ggplot2 (Wickham, 2016), Pheatmap (Kolde, 2019) and plotly (Plotly Technologies Inc, 2015).

### Web application development

We built a web application using ‘Shiny’ with the shinydashboard template (Chang *et al*., 2017; Chang and Ribeiro, 2018). The gene expression database used in the application is stored in a partitioned parquet directory. The interface between R and the Apache Arrow platform is performed with the ‘Arrow’ R package (Richardson *et al*., 2021). The app is freely accessible at https://cannatlas.venanciogroup.uenf.br/.

## 3 Resource overview

We developed the Cannabis Expression Atlas, a web application designed to explore the expression patterns of Cannabis genes using RNA-Seq data. This atlas currently comprises the expression of 27,640 genes across 394 RNA-seq samples, along with sample metadata (Table S1). It consists of four main pages, each offering different data navigation options (Figure 1):

**Figure 1.**
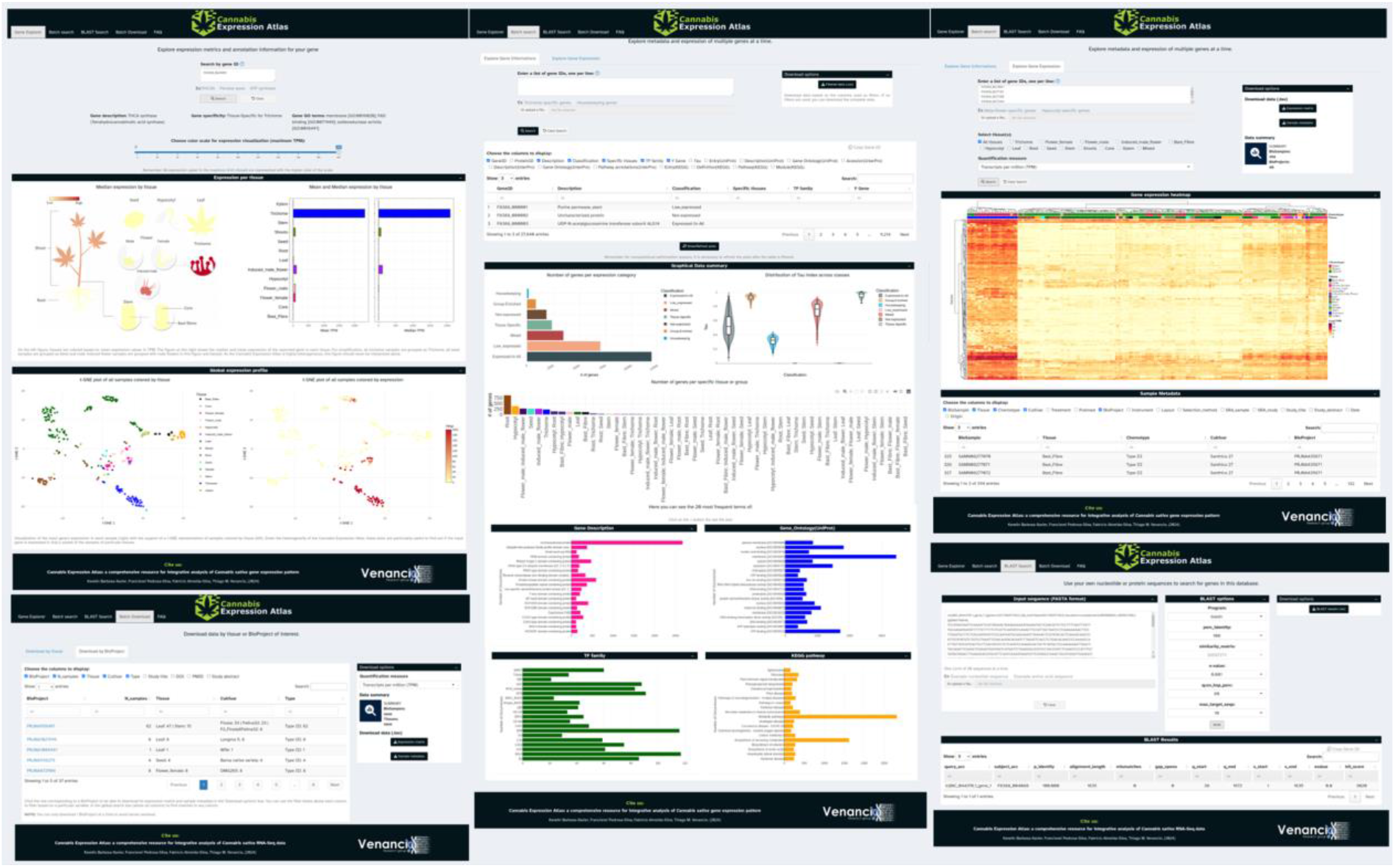
Overview of the Cannabis Expression Atlas.

I. **Gene explorer:** users can search individual Gene IDs and explore their expression profiles through t-SNE, median, and mean TPM barplots. A representative image of the tissues colored according to the median expression levels of the gene of interest is also provided.
II. **Batch search:** users can fetch the expression of multiple genes. In this case, there are two navigation options:
  A. **Gene information tab:** users can submit gene lists and explore a dynamic table that can be filtered by Gene ID, classification, tissue, GO terms, pathways, among other annotation information. Different plots, statistics, and download options are also provided.
  B. **Gene expression tab:** users can submit a list of genes and explore their expression across tissues, represented in a heatmap that is generated along an accessory table with sample metadata.
III. **BLAST Search:** users can find genes of interest by querying nucleotide or amino acid sequences.
IV. **Batch download:** in this page, users can fetch expression data by one of two tabs:
  A. **Download by tissue:** selection of the tissue of interest and quantification method (TPM or counts);
  B. **Download by BioProject:** users can filter a table by cultivar, chemotype, or publication keywords, DOI, PMID, and title. The web app then generates an expression matrix and a file with the relevant metadata.

## 4 Results and discussion

### Publicly available Cannabis RNA-seq data comprise 394 high-quality samples of 13 tissues, 55 varieties, and three chemotypes

In August 2024, we identified 515 publicly available RNA-Seq samples from the SRA database. Following download and preprocessing, which included read trimming to remove adapters and low-quality bases, we excluded 40 samples that exhibited a mean length of less than 40 bp or a Q20 rate below 80% (Figure S1a, b, c). These samples belong to only four bioprojects (Table S4a). We also removed 81 samples with read mapping rates lower than 50% (Figure S1a, d; Table S4b), resulting in a database of 394 high-quality samples. By retaining only high-quality samples, we improve the reliability of gene expression quantification and minimize the prevalence of potentially contaminated samples (Almeida-Silva, Pedrosa-Silva and Venancio, 2023). By using this filtered dataset, we successfully quantified transcript abundance of 27,640 Cannabis genes representing 99% of the reference transcriptome (JL (27,358) + Y genes (535) = 27,893 genes).

The atlas encompasses 13 different plant tissues and the mixed samples (Figure 2a, c), with leaf representing 38.3% of the samples. We grouped these 394 samples into the three main chemotypes (Figure 2b), namely type I (marijuana), type II (balanced), and type III (hemp). Furthermore, these samples originate from 55 Cannabis varieties with the four most represented being Finola (n = 42), Santhica 27 (n = 36), a F1 from Agp029 x blackberry kush auto (n = 27), and Felina32 (n = 23) (Figure 2d), of which the F1 from Agp029 x blackberry kush is a type I and the other three are type III. The summary statistics reveal that 76.5% of the Cannabis RNA-seq data available in public databases are of good quality, with the mapping rate filter being responsible for the exclusion of 15.7% of the samples (Figure S1a, d). Most samples demonstrated high mapping rates, with an average of approximately 80% and a smaller cluster around 40% (Figure S1d). The median and mean number of reads per sample were calculated at 42 and 51 million reads, respectively (Figure S1b). Paired-end sequencing was used for most samples (366), while the Illumina NovaSeq 6000 platform was employed for 153 samples.

**Figure 2.**
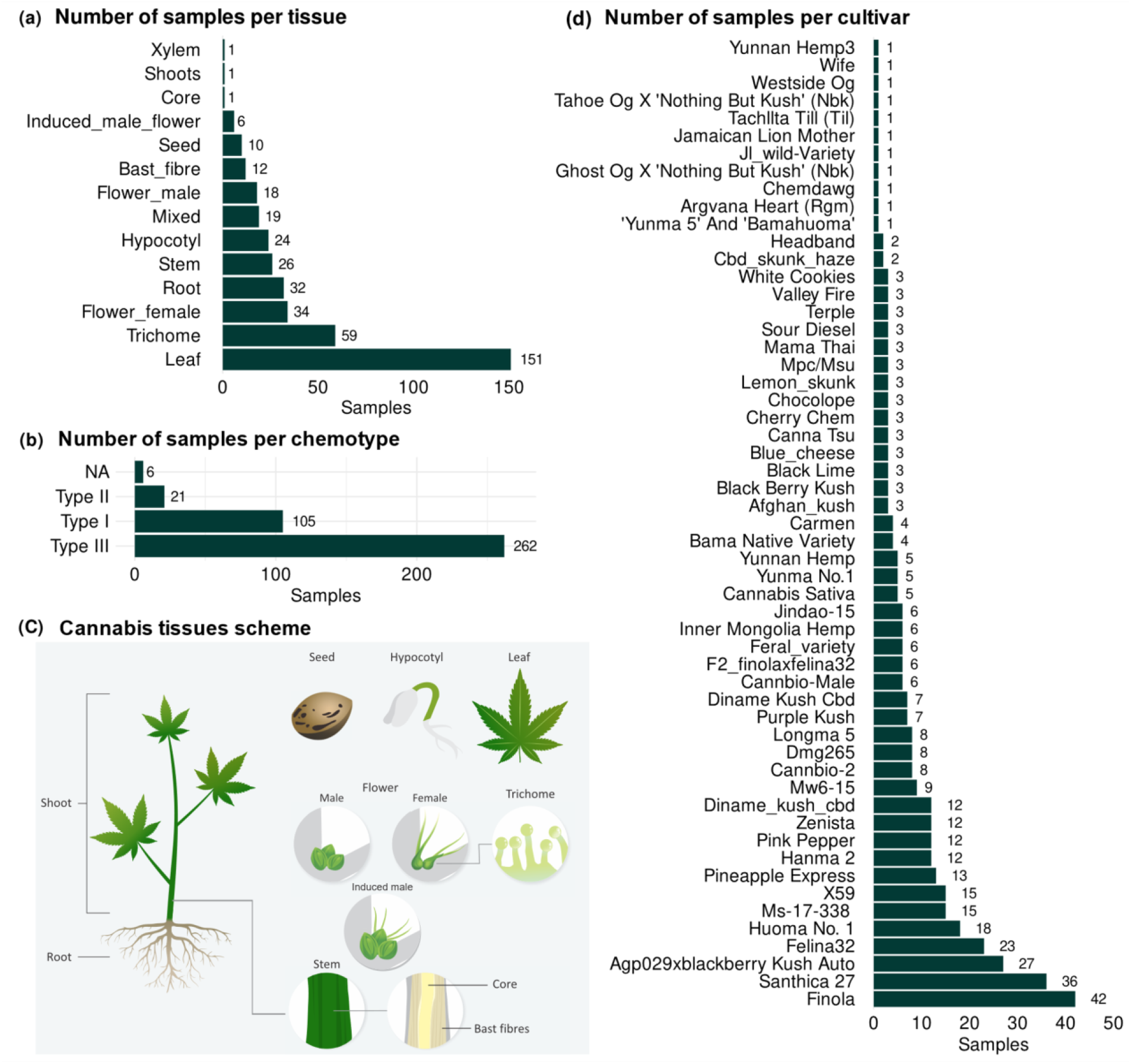
Distribution of Cannabis samples. (a) Number of samples per tissue. (b) Number of samples per chemotype. (c) Schematic representation of Cannabis tissues. (d) Number of samples per cultivar.

### The increase in RNA-Seq data highlights the growing interest in Cannabis research

The first publication of a Cannabis RNA-seq study dates back to 2011 (Bakel *et al*., 2011). However, a significant number of studies emerged only after 2016 (Figure S1e, f, g), coinciding with the global legislative shift on Cannabis use and research (United Nations, 2024). Notably, type III (hemp) samples outnumbered type I (marijuana) and type II (balanced) samples (Figure S1f), reflecting industrial and agricultural interests. Interestingly, since 2019, there has been a notable increase in the number of samples derived from plant tissues associated with medicinal applications, such as trichomes and female flowers, which correlates with an increase in types I and II samples (Figure S1f, g) which highlights the growing emphasis on the Cannabis therapeutic potential. Over 51% of the samples were contributed by research groups based in China and Canada (Figure S1h). These RNA-Seq studies addressed a myriad of Cannabis research questions, including fiber production and quality (Guerriero *et al*., 2017), biotic and abiotic resistance (Gao *et al*., 2018; McKernan *et al*., 2020; Cao *et al*., 2021, 2023; Jiang *et al*., 2021; Pépin, Hebert and Joly, 2021; Yin *et al*., 2022; Yan *et al*., 2023), sex determination (Prentout *et al*., 2020; Adal *et al*., 2021; Dowling *et al*., 2023), metabolite production, quality, chemotype identification (Braich *et al*., 2019; Laverty *et al*., 2019; Zager *et al*., 2019; Booth *et al*., 2020; Livingston *et al*., 2020; McGarvey *et al*., 2020; McKernan *et al*., 2020; Busta *et al*., 2022; Yeo *et al*., 2022; Mi *et al*., 2023; Tang *et al*., 2023), and genome assembly (Bakel *et al*., 2011; Braich *et al*., 2020; Gao *et al*., 2020; McKernan *et al*., 2020).

### Dimensionality reduction reveals distinct transcriptional profiles by tissue and chemotype

The elbow point statistics indicated that 8 principal components account for 65% of the expression variation. The t-SNE dimensionality reduction using these components shows that the most significant variation associated with chemotype was observed in leaves, where two distinct clusters of type I samples were separated from type III samples (Figure 3). The two Type I leaf clusters are of two different varieties and BioProjects, with one (the bigger leaf type I cluster, Figure 3) being focused on resistance to *Golovinomyces ambrosiae* (PRJNA738505) and the other (the smallest leaf type I cluster, Figure 3) focused on study auto-flowering (PRJNA1049889). Fiber-related tissues, such as hypocotyl, bast fiber, and stem formed a tightly grouped cluster, reflecting their shared role in structural support and potential similarity in gene expression related to cell wall biosynthesis and secondary growth. Reproductive tissue samples (female, male and induced male flowers) clustered closely although not very cohesively (Figure 3) and a divergence between female and male flowers is noticeable. Chemotype and maturity associated clustering patterns were also observed. Trichomes exhibited comparable global expression patterns regardless of chemotype. Finally, root samples were clearly distinct from all other tissue types.

**Figure 3.**
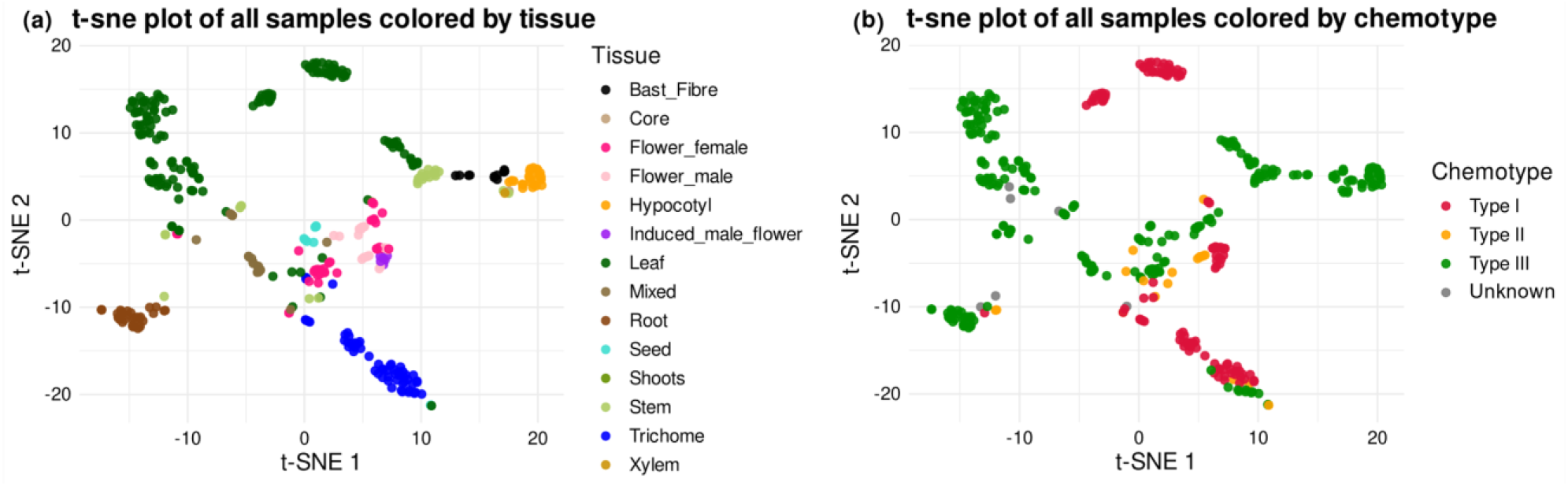
Dimensionality reduction performed with t-SNE. (a) Samples colored by tissue. (b) Samples colored by chemotype.

### The Cannabis Expression Atlas comprises seven gene expression categories and over 1400 transcription factors

We classified genes into seven expression categories (Figure 4, Table S2) (Jain and Tuteja, 2019; Machado *et al*., 2020; Almeida-Silva, Pedrosa-Silva and Venancio, 2023). Housekeeping genes (HK, 132 genes) were identified as those with consistently high expression and low variation across samples (Figure S2a) which support their roles in fundamental cellular functions and physiological processes (Joshi *et al*., 2022). Tissue-specific genes (2,382 genes) were defined by a TAU index between 0.8 and 1.0 and at least five-fold higher expression in one tissue compared to others (Figure 4 and Figure S2b). These genes can shed light on the specific regulatory mechanisms governing the development of various plant parts. Group-enriched genes (843 genes) have at least five-fold higher expression levels in a group of 2 tissues in comparison to all other tissues. The Expressed-in-all category comprises 11,929 genes expressed in all tissues. The Low-expressed group (7,021 genes) had TPM ≥ 1 in at least one sample but median TPM ≤ 5, while Not-expressed genes (1,853) had TPM < 1 in all samples. Finally, 3,480 genes were classified as mixed, not fitting into the previous categories.

**Figure 4.**
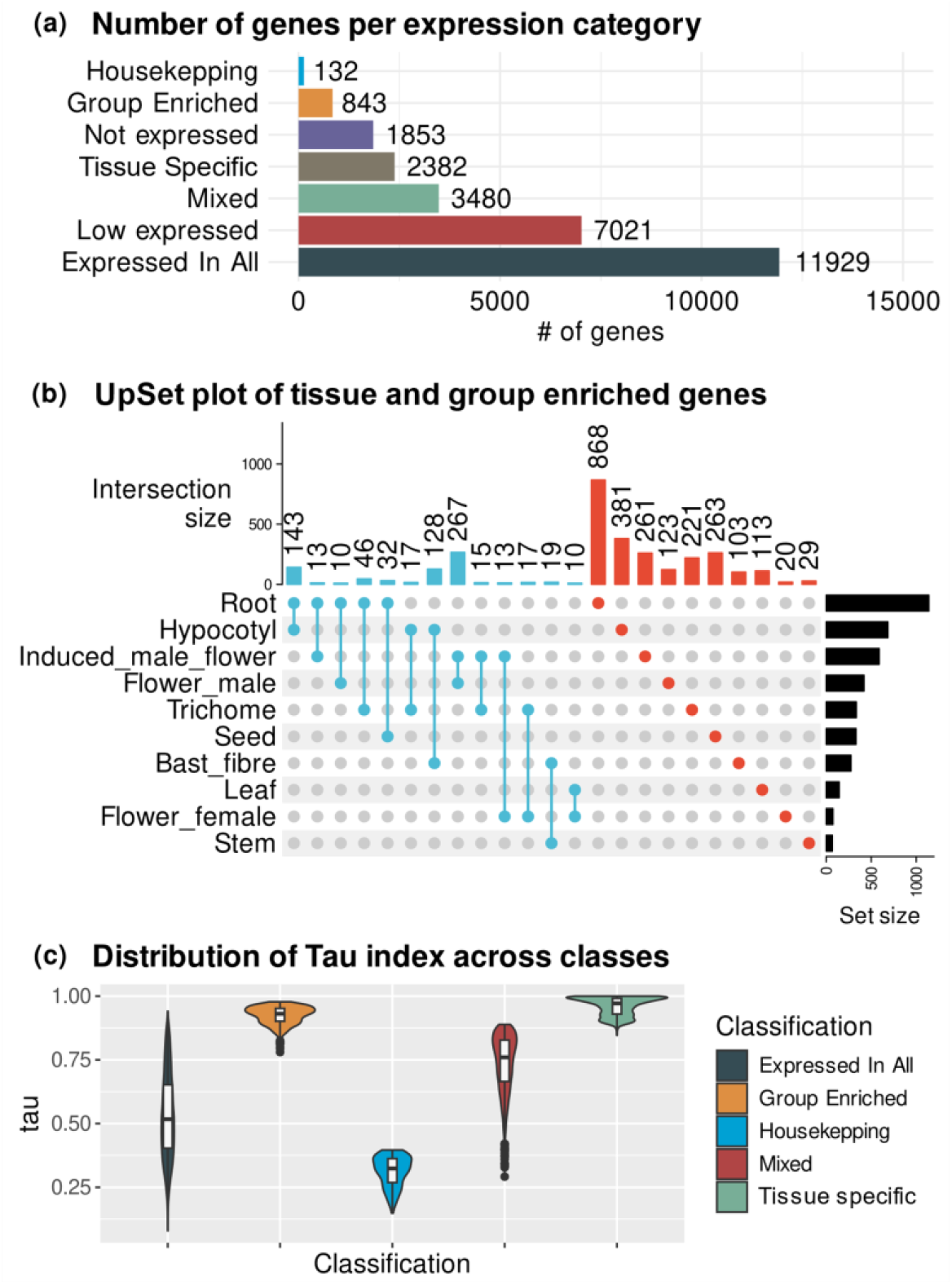
Gene expression classification. (a) Number of genes per expression category. (b) UpSet plot of tissue and group specific genes. (c) Violin plot of distribution of the Tau index across gene classes.

By utilizing the reference genome (see methods for details), we identified 535 Y-linked genes of the JL father, termed Y-genes for brevity. Of these, 370 had detectable expression in our dataset (see methods for details), out of which 130 showed median TPM ≥ 5 and were analyzed for tissue-specific expression. From Y-genes, 240 were Low-expressed, 165 were Not-expressed, 61 were specifically expressed in male flowers, 26 showed mixed expression, 22 were expressed-in-all and 11 were group-enriched (Table S2). The Y-genes classified as mixed or expressed-in-all suggest that they are located in the pseudoautosomal region of Y chromosome (Divashuk *et al*., 2014). This list of Y-genes and their expression patterns can offer insights into sex determination and male flower development and physiology.

Out of the 27,640 genes in the Cannabis Expression Atlas, we found 1,489 TFs using PlantTFDB (Table S2) as a reference, out of which 177 (11.8%) were tissue-specific (Table 1). The top 10 TF families in number of genes were bHLH, ERF, MYB, NAC, B3, MYB_related, C2H2, bZIP, WRKY, and C3H (Table S3) and correspond to 56.1% of all TFs of the Cannabis Expression Atlas. The highest prevalence of tissue specific TFs was observed in roots (n = 75), followed by hypocotyl (n = 30), seed (n = 21), and induced male flowers (n = 17) (Table 1). This pattern aligns with the high number of specific genes identified in these tissues (Figure 4b). These TFs can be important regulators of tissue-specific regulatory mechanisms and constitute important candidates for biotechnological applications.

**Table 1.**
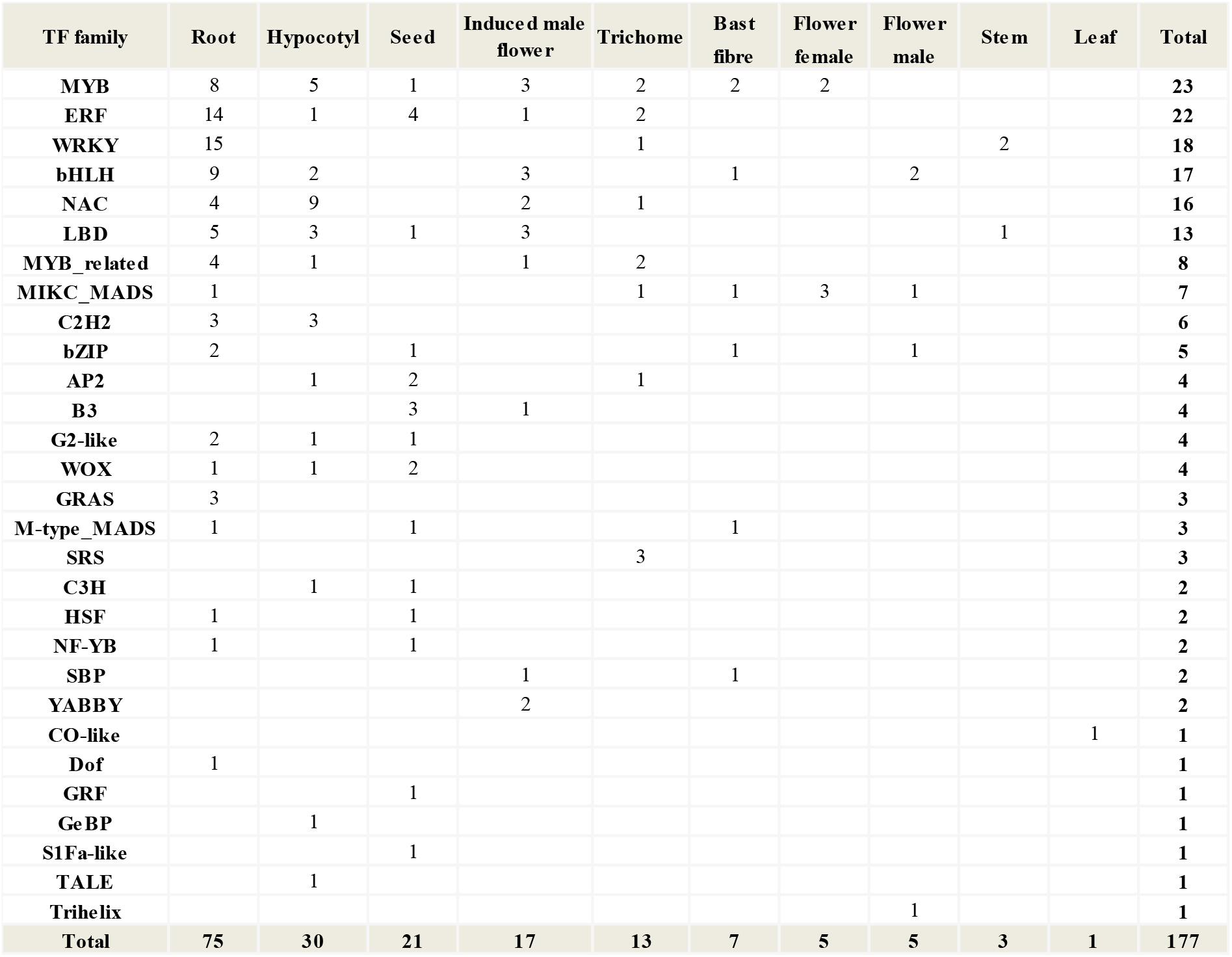
Tissue-specific transcription factors.

Interestingly, the Cannabis Expression Atlas has more TFs than that reported in PlantTFDB 5.0 for Cannabis (Tian *et al*., 2020). Table 2 presents the top 10 TF families in the Cannabis Expression Atlas. This discrepancy may be attributed to the fact that the Cannabis TFs in PlantTFDB 5.0 are based on the Purple Kush genome, which has just 78.1% of BUSCO completeness score, whereas the one used here achieved 96.8% (Medicinal Genomics, 2019). The progress in genome sequencing and assembly likely impacted gene predictions.

**Table 2.** Top 10 transcription factors family, in number of genes, at Cannabis Expression Atlas compared to PlanTFDB.

| TF_family | Cannabis Expression Atlas members | PlantTFDB Cannabis Members (v5.0) | Fold change (Cannatlas/PlantTFDB) |
| --- | --- | --- | --- |
| bHLH | 117 | 99 | 1.18 |
| ERF | 116 | 59 | 1.96 |
| MYB | 91 | 81 | 1.12 |
| NAC | 88 | 75 | 1.17 |
| B3 | 86 | 60 | 1.43 |
| MYB_related | 83 | 70 | 1.18 |
| C2H2 | 75 | 62 | 1.20 |
| bZIP | 61 | 54 | 1.12 |
| WRKY | 60 | 49 | 1.22 |
| C3H | 59 | 48 | 1.22 |
| Total | 836 | 657 | 1.27 |

## 5 Conclusions

The development of the Cannabis Expression Atlas represents significant progress in the field, providing researchers with a user-friendly and freely available platform to explore gene expression data. This tool facilitates the investigation of gene expression patterns across different tissues and conditions, enabling researchers to conduct in-depth analyses and generate new hypotheses. The integration of metadata and various search options enhances the usability of the atlas, making it a valuable resource for the research community.

## Author contributions

Conceived the study: K.B.-X. and T.M.V.; Funding and resources: T.M.V.; Data analysis: K.B.-X.; Interpretation of the results: K.B.-X. and T.M.V.; Source development: K.B.-X., F.A.-S., F.P.-S. and T.M.V.; Wrote the manuscript: K.B.-X. and T.M.V.

## Acknowledgment

This work was supported by Fundação Carlos Chagas Filho de Amparo à Pesquisa do Estado do Rio de Janeiro (FAPERJ), Coordenação de Aperfeiçoamento de Pessoal de Nível Superior-Brasil (CAPES; Finance Code 001), and Conselho Nacional de Desenvolvimento Científico e Tecnológico CNPq). The funding agencies had no role in the design of the study and collection, analysis and interpretation of data and in writing.

## Data availability and FAIR (Findable Accessible Interoperable Reusable) compliance statement

The Cannabis Expression Atlas container is available at [Docker hub](https://hub.docker.com/r/barbosaxavierkevelin/cannabisexpressionatlas) and the parquet directory is available at [FigShare](https://figshare.com/s/f5fe2fd793bd4f74949d). All the data files, supplementary materials and methodology code used here are available at [GitHub](https://github.com/barbosaxavierkevelin/CannAtlas-paper.git).

