## Supplementary figures for "Cannabis Expression Atlas: a comprehensive resource for integrative analysis of *Cannabis sativa* L. gene expression"

[1] Laboratório de Química e Função de Proteínas e Peptídeos, Centro de Biociências e Biotecnologia, Universidade Estadual do Norte Fluminense Darcy Ribeiro, Campos dos Goytacazes, RJ, Brazil.

[2] Department of Plant Biotechnology and Bioinformatics, Ghent University, Ghent, Belgium.

[3] VIB Center for Plant Systems Biology, VIB, Ghent, Belgium.

* TMV: Laboratório de Química e Função de Proteínas e Peptídeos, Centro de Biociências e Biotecnologia, Universidade Estadual do Norte Fluminense Darcy Ribeiro. Av. Alberto Lamego 2000, P5, sala 217, Campos dos Goytacazes, RJ, Brazil..

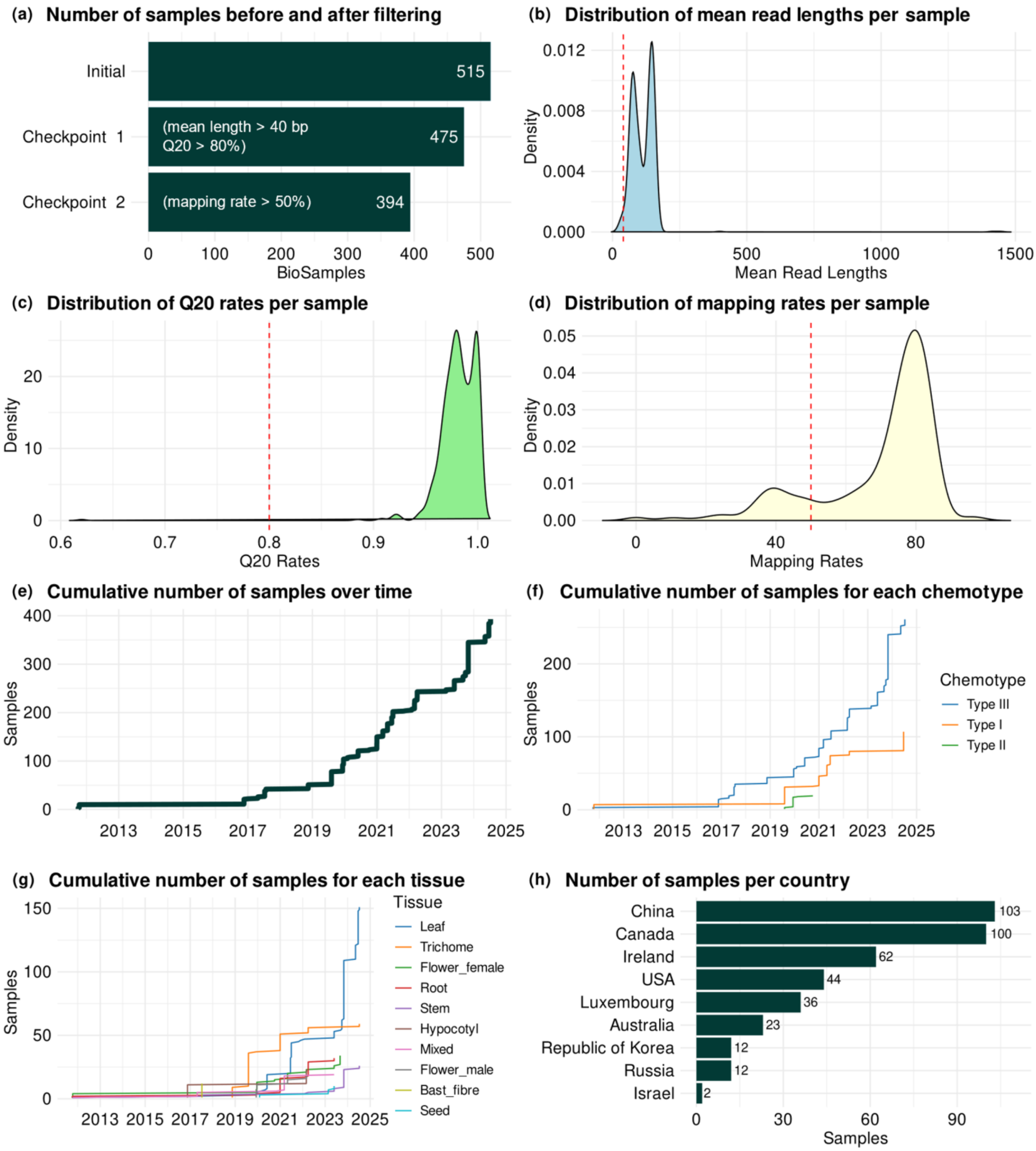


**Figure S1.** Data summary quality statistics and evolution of the number of samples deposited in NCBI SRA.. (a) Number of samples before and after filtering steps. (b) Distribution of mean read lengths per sample. Dashed red line represents the minimum read length threshold from checkpoint 1. (c) Distribution of Q20 rates per sample. Dashed red line represents the minimum Q20 rate threshold from checkpoint 1. (d) Distribution of mapping rates per sample. Dashed red line indicates the minimum mapping rate threshold from checkpoint 2. (e) Cumulative number of samples over years. (f) Cumulative number of samples by chemotype over years. (g) Cumulative number of samples by tissue over years. Tissues with fewer than 10 samples are not displayed. (h) Number of samples by country. Samples removed during filtering steps are not considered in these panels.

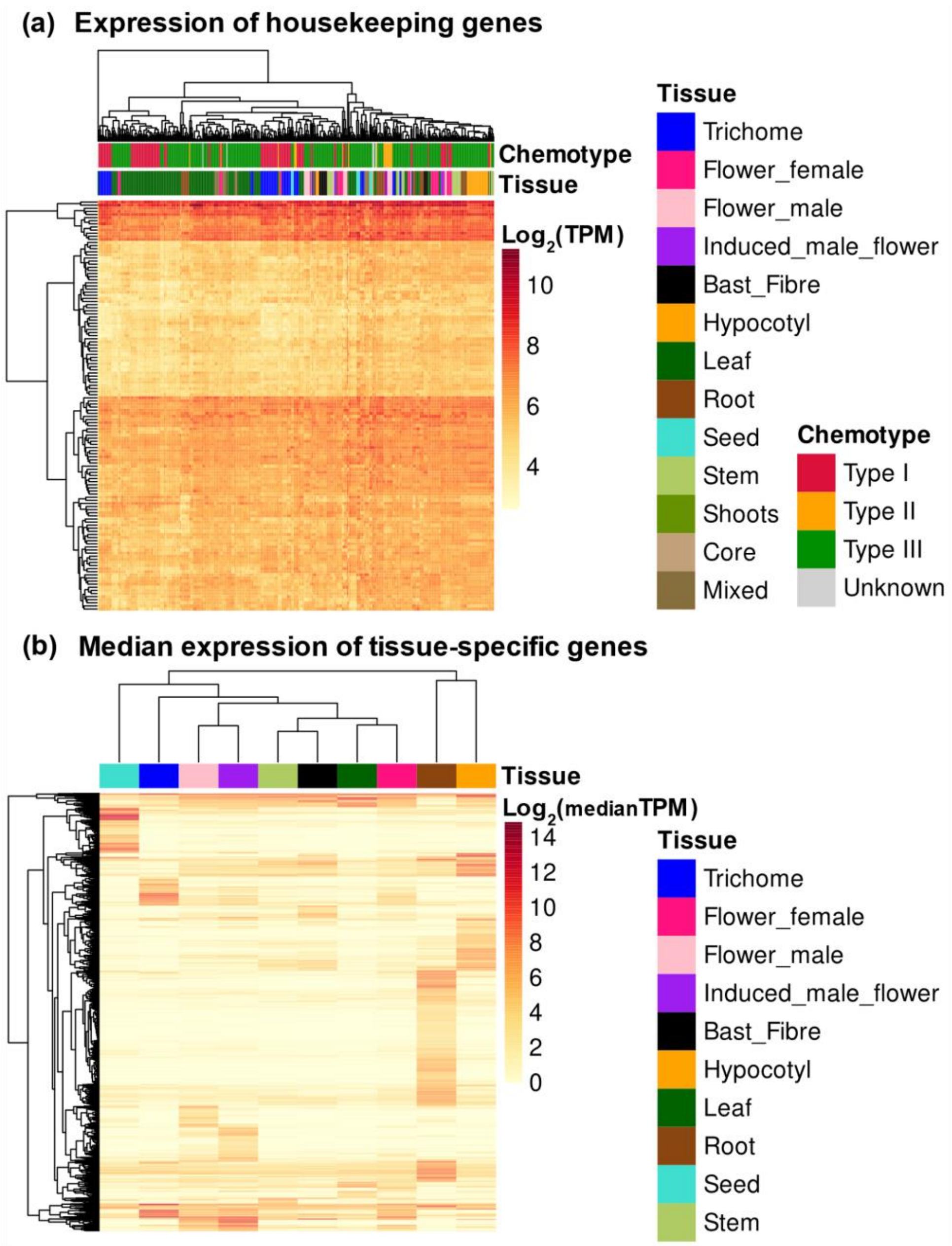


**Figure S2.** Heatmap of (a) housekeeping genes expression in log2(TPM) and (b) tissue-specific genes median expression by tissue in log2(median(TPM)).
